# Systematic identification of cis-interacting lncRNAs and their targets

**DOI:** 10.1101/2021.01.13.426305

**Authors:** Saumya Agrawal, Ivan V. Kulakovskiy, Jessica Severin, Masaru Koido, Tanvir Alam, Imad Abugessaisa, Andrey Buyan, Howard Y. Chang, Josee Dostie, Masayoshi Itoh, Juha Kere, Naoto Kondo, Yunjing Li, Vsevolod J. Makeev, Mickaël Mendez, Yasushi Okazaki, Jordan A. Ramilowski, Andrey I. Sigorskikh, Lisa J. Strug, Ken Yagi, Kayoko Yasuzawa, Chi Wai Yip, Chung Chau Hon, Michael M. Hoffman, Chikashi Terao, Takeya Kasukawa, Jay W. Shin, Piero Carninci, Michiel JL de Hoon

**Affiliations:** RIKEN Center for Integrative Medical Sciences, Yokohama, Japan; Institute of Medical Science, The University of Tokyo, Tokyo, Japan; College of Science and Engineering, Hamad Bin Khalifa University, Doha, Qatar; Center for Personal Dynamic Regulome, Stanford University, Stanford, California, USA; Department of Biochemistry, Rosalind and Morris Goodman Cancer Research Center, McGill University, Montréal, Québec, Canada; RIKEN Preventive Medicine and Diagnosis Innovation Program, Wako, Japan; Department of Biosciences and Nutrition, Karolinska Institutet, Huddinge, Sweden; Stem Cells and Metabolism Research Program, University of Helsinki and Folkhälsan Research Center, Helsinki, Finland; RIKEN Center for Life Science Technologies, Yokohama, Japan; Division of Biostatistics, Dalla Lana School of Public Health, University of Toronto, Toronto, Ontario, Canada; Department of Computer Science, University of Toronto, Toronto, Ontario, Canada; Advanced Medical Research Center, Yokohama City University, Yokohama, Japan; Division of Biostatistics, University of Toronto, Ontario, Canada; The Centre for Applied Genomics and Program in Genetics and Genome Biology, The Hospital for Sick Children, Toronto, Ontario, Canada; Princess Margaret Cancer Centre, Toronto, Ontario, Canada; Department of Medical Biophysics, University of Toronto, Toronto, Ontario, Canada; Vector Institute, Toronto, Ontario, Canada; Human Technopole, Milan, Italy; autosome.org; FANTOM Consortium

**Author notes:** These authors contributed equally to this work (Ivan V. Kulakovskiy, Jessica Severin, Masaru Koido, Tanvir Alam).

## Abstract

The human genome is pervasively transcribed and produces a wide variety of long non-coding RNAs (lncRNAs), constituting the majority of transcripts across human cell types. Studying lncRNAs is challenging due to their low expression level, cell type-specific occurrence, poor sequence conservation between orthologs, and lack of information about RNA domains. LncRNAs direct the regulatory factors in the locations that are in cis to their transcription sites. We designed a model to predict if an lncRNA acts in cis based on its features and trained it using RNA-chromatin interaction data. The trained model is cell type-independent and does not require RNA-chromatin data. Combining RNA-chromatin and Hi-C data, we showed that lncRNA-chromatin binding sites are determined by chromosome conformation. For each lncRNA, the spatially proximal genes were identified as their potential targets by combining Hi-C and Cap Analysis Gene Expression (CAGE) data in 18 human cell types. RNA-protein and RNA-chromatin interaction data suggested that lncRNAs act as scaffolds to recruit regulatory proteins to target promoters and enhancers. We provide the data through an interactive visualization web portal at https://fantom.gsc.riken.jp/zenbu/reports/#F6_3D_lncRNA.

## Background

Human cells express tens of thousands of long non-coding RNAs (lncRNAs)[1], defined as RNA transcripts of at least 200 nt with no or limited protein-coding potential. Although this class of RNAs has been known for almost 50 years [2–4], ∼95% of lncRNAs still lack functional annotation [5], though some lncRNAs have been shown to have important roles in transcriptional regulation [6, 7], chromatin maintenance [8, 9], translation [10], and other biological processes. Exploring the role of the lncRNAs systematically is challenging due to their low expression [11], rapid degradation compared to mRNAs [12, 13], high cell type-specificity [11], and lack of conservation across organisms [14]. In contrast to protein-coding genes, the absence of families of lncRNAs with related sequences further hinders the systematic inference of lncRNA target regions.

Even in low copy numbers, lncRNAs can initiate the formation of nuclear compartments by interacting with RBPs to regulate transcription [15]. LncRNAs are reported to form scaffolds with RBPs and other mediator proteins that regulate the process of transcription and chromatin remodeling [16–18]. Studies using microscopy and RNA-chromatin interaction sequencing techniques have shown that the majority of lncRNAs are located in the nucleus and are likely constrained to regions close to their transcription site in three-dimensional space [19, 20]. In the majority, lncRNAs are more stable than nascent mRNA transcripts hence these signals are unlikely to be due to the act of transcription of lncRNAs but to stable interactions [12]. The lncRNA’s regulatory role may be associated with the transcript itself or the act of transcription of its promoter [21]. Regardless of the mode of action, lncRNAs can regulate the expression of genes several megabases away in linear genomic distance [22–24], [25, 26] as distal regions are brought into spatial proximity by chromatin folding.

LncRNA retention in nucleus is directed by the presence of different RNA features [27, 28] including repeat sequences [29], inverted Alu elements [30, 31], motifs like SIRLOIN [32] and U1-recognition motif [33]. These features were combined with nuclear/cytoplasmic fraction and RNA-chromatin interaction data to predict the likelihood of each lncRNA (which we termed as **cis-score**) in different cell types to interact with chromatin in cis (with regions that are on the same chromosome as the lncRNA promoter). As a proof of principle, using RNA-chromatin interaction data we show that by combining the chromatin interaction data (Hi-C) with the cis-score we can identify potential interacting partners of a lncRNA. High-resolution genomic interaction maps were generated using newly generated Hi-C data for induced pluripotent stem cells (iPSCs) and previously published data from 17 other human cell types and tissues. These interaction maps were employed to systematically identify candidate targets of lncRNAs. Cap Analysis Gene Expression (CAGE) [34] data was used to determined expressed genes and their precise transcription start sites (TSSs), and bi-directionally transcribed enhancers. With these data, we identified lncRNA interacting partners (including mRNAs, enhancers, and other lncRNAs) genes as candidate targets of lncRNAs across the 18 cell types and tissues. A visualization platform is provided that allows users to browse and compare the biological features for each lncRNA in individual cell types (https://fantom.gsc.riken.jp/zenbu/reports/#F6_3D_lncRNA).

## Results

### Candidate cis-acting lncRNAs

To construct the model to determine the putative cis-acting lncRNAs, we integrated information regarding the fraction of lncRNAs in the nucleus compared to the cytoplasm, expression level, frequency of repeat sequences [29], presence of inverted Alu elements in exons [30, 31], SIRLOIN motif [32] and U1-recognition motif [33] together with RNA-chromatin interactions in seven different human cell types (embryonic cell type: iPSC, primary cell type: HFFc6, and HUVEC, cell line: HEK293T, MDA231, K562, and MM1S [35–37]; [38]; [39, 40]). LncRNAs with ≥ 75% of raw RNA-chromatin interactions in cis are significantly more likely to have these sequence elements compared to remaining lncRNAs **(Figure 1; Table-1)**. We predict the probability of a lncRNA to act in cis by combining all six factors. The contribution of each element towards the cis score increases in the order: inverted Alu repeat < SIRLOIN < repeat elements < U-repeat motif ≤ abs[log10(expression level)] < nuclear percentage (**Table-2**). Since iPSC has the deepest sequenced RNA-chromatin interaction data therefore the model calculated using its data was used to estimate the score for all the selected cell types **(Table-3)**. To benchmark, if the iPSC model is optimum, we compare the scores predicted using iPSC cells and models generated using the RNA-chromatin data from the same cell type. The sensitivity to predict if lncRNA potentially acts in cis increased while specificity decreased when using the iPSC model compared to the same cell model **(Supplementary Figure 1)**. This was expected, as compared to iPSC the sequencing depth for all the other cell types is shallow therefore, is likely to miss several real RNA-chromatin interactions. Next, to verify that the cis-score does represent lncRNA localization, we compared the cis-scores for lncRNAs to the localization patterns determined using single-molecule microscopy [19]. For the lncRNAs where the signal was punctate, the scores are higher regardless of cell types while lncRNAs with dispersed signals show variation in cis-score (**Table-4**). Overall, this supports that the calculated scores can predict the localization of lncRNA in the nucleus.

**Figure 1:**
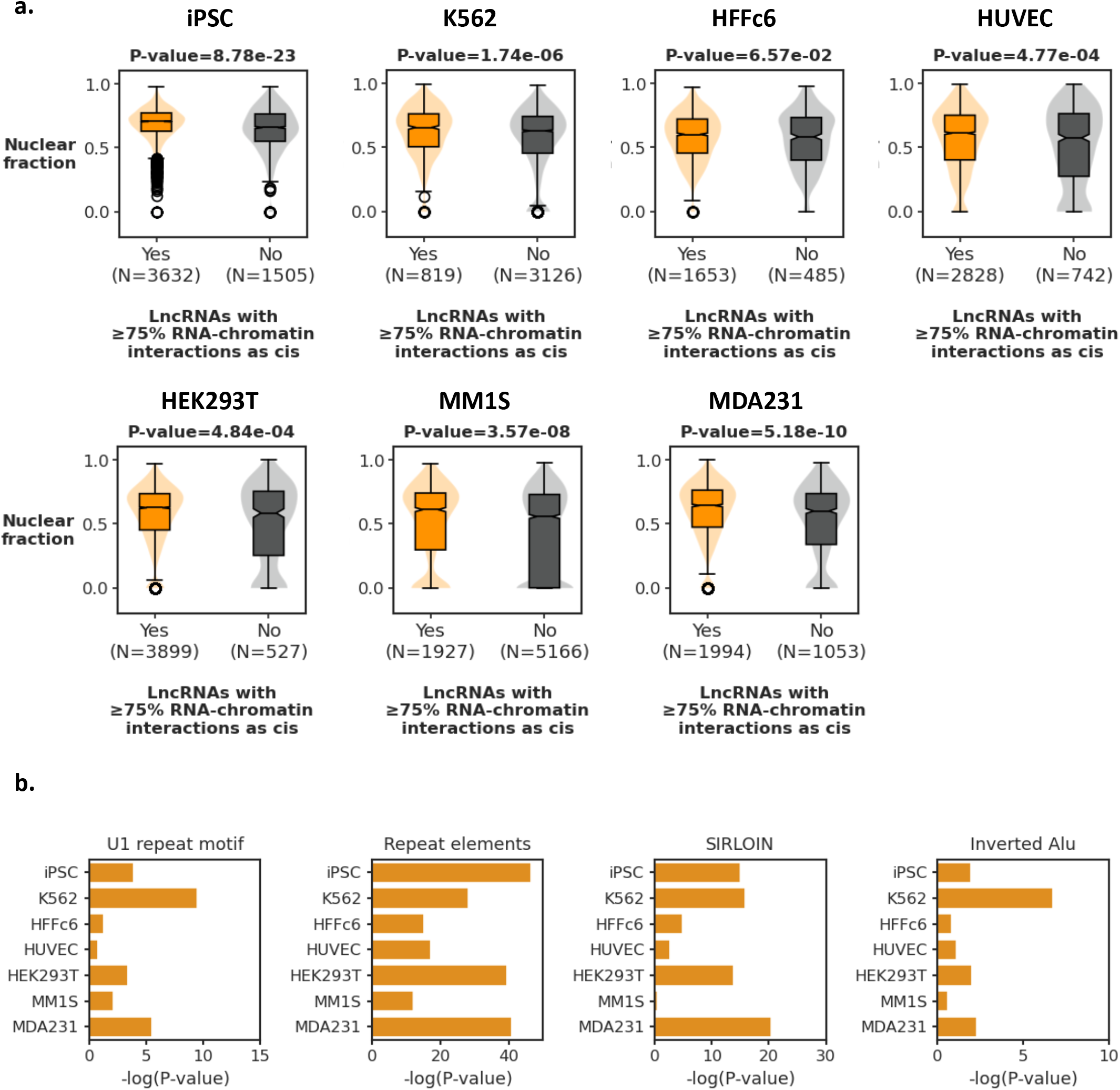
Comparison of cis-score parameters between lncRNAs with ⩾75% of raw RNA-chromatin interaction in cis and less. **a)** Fraction of lncRNA in the nucleus compared to the cytoplasm. The significance of the difference in values between two classes of lncRNAs was determined using the one-tailed Mann-Whitney U test. **b)** Statistical significance of enrichment of lncRNAs with RNA features: U1 repeat motif count, frequency of repeat sequences, SIRLION motif count, and inverted Alu elements, in the lncRNAs with ⩾75% RNA-chromatin interaction are cis. Significance was calculated using a one-tailed Fisher’s exact test.

### The target regions of lncRNAs are in close proximity

Raw RNA-chromatin interaction data show that lncRNAs with higher cis-scores tend to interact with more genomic regions (rho = 0.3-0.5; **Supplementary Figure 2**). Comparison of RNA-chromatin and Hi-C interactions data from the same cell type (**Table 5-7**) shows that the captured raw RNA-chromatin interactions formed in the regions that are linearly several megabases away from the promoter window of the lncRNA but are in close proximity in physical space as shown by the Hi-C data **(Figure 2a)**. Further, this linear distance for captured RNA-chromatin interaction increases with an increase in the half-life of the lncRNAs **(Supplementary Figure 3)**. Overall, the majority of lncRNA interactions are directed by the chromatin conformation of the region and lncRNA stability. The cis-score of the lncRNAs with atleast one significant RNA-chromatin interaction (FDR ≤ 0.01) is significantly higher compared to ones that have none **(Figure 2b; Supplementary Figure 4; Supplement data 1)**. Further, based on Hi-C interactions, on average 87.5% of the genomic regions to which an RNA binds are within 2 degrees of separation from its promoter (**Figure 2c**) and are in the same A/B compartment as the gene encoding the RNA **(Table 8)**. All together, this shows that for the majority of lncRNAs, combining cis-score with Hi-C can identify the target regions detected by RNA-chromatin capture technologies.

**Figure 2:**
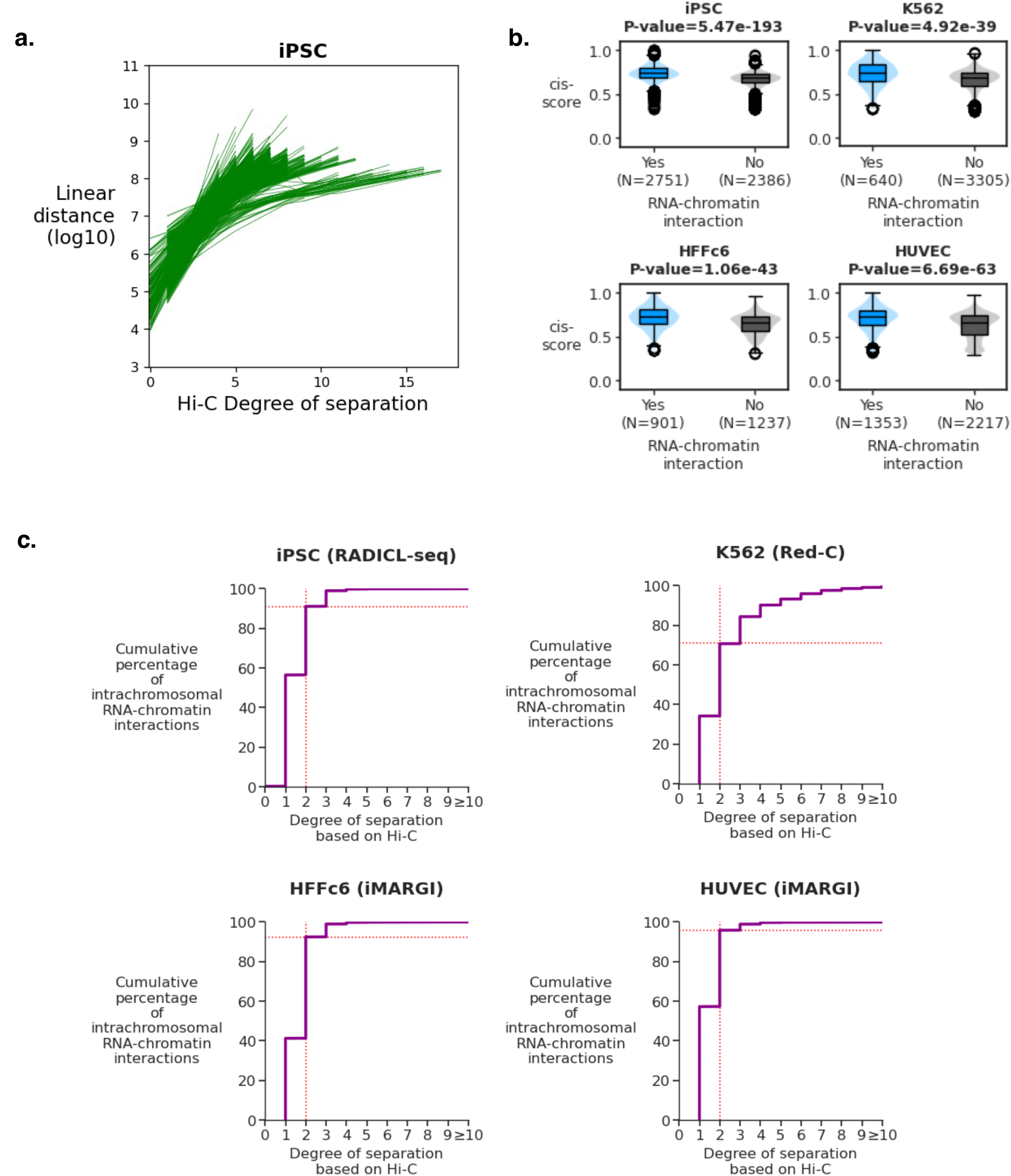
The comparison between RNA-chromatin interactions and Hi-C interactions. **a**. Hi-C degree of separation vs Linear distance for lncRNA raw RNA-chromatin interaction in iPSC using local regression. The horizontal axis and vertical axis show the degree of separation (calculated using Hi-C interactions) and linear distance between the windows with the promoter of the lncRNA and the region containing RNA-chromatin interaction. Each line represents one lncRNA. **b.** Comparison of cis-score between lncRNAs with and without atleast one RNA-chromatin interaction. Each panel shows one cell type for which RNA-chromatin data is available. The significance of difference in cis-score between two classes of lncRNAs was determined using a one-tailed Mann-Whitney U test. **c.** Cumulative distribution plot showing degree of separation between chromatin regions where RNAs bind based on Hi-C interactions.

### Defining candidate target genes for each lncRNA and their biological features in a different cell type

To increase the scope of the study we have further processed previously published Hi-C data for one embryonic cell line, four primary cell types, eight cell lines, and one tissue **(Table 5-7)** were uniformly processed. As outlined in **Figure 3,** for every expressed lncRNA in each cell type, the genomic regions were selected with up to 2 degrees of separation (cutoff decided based on RNA-chromatin and Hi-C interaction comparison) from the region containing the reference lncRNA gene. Next, promoters and enhancers expressed in the cell type were mapped to the genomic regions to identify the candidate targets for every lncRNA in each cell type. Gene Ontology (GO) enrichment analysis for the candidate target genes of every lncRNA independently in each cell type showed that the predicted target genes of several lncRNAs had at least one significantly enriched “biological processes” GO term **(Table 9; Table 10)** suggesting that certain lncRNAs can coordinate multiple related genes. The lncRNAs facilitate the formation of condensates which are thought to assist in the recruitment of the transcription factors [41, 42]. The transcription factor binding sites (TFBSs) enrichment analysis for the candidate target genes and enhancers of each lncRNA showed that multiple lncRNA’s candidate targets were significantly enriched for at least one motif **(Table 11-12)**. Further, lncRNA candidate target genes showed a significant enrichment in genes with promoters with either a high or a low GC content **(Supplementary Figure 5, Table 13)**, which has a decisive role in determining transcription factor binding [43–46] and may contribute to the TFBS enrichment observed for the candidate target genes of each lncRNA.

**Figure 3:**
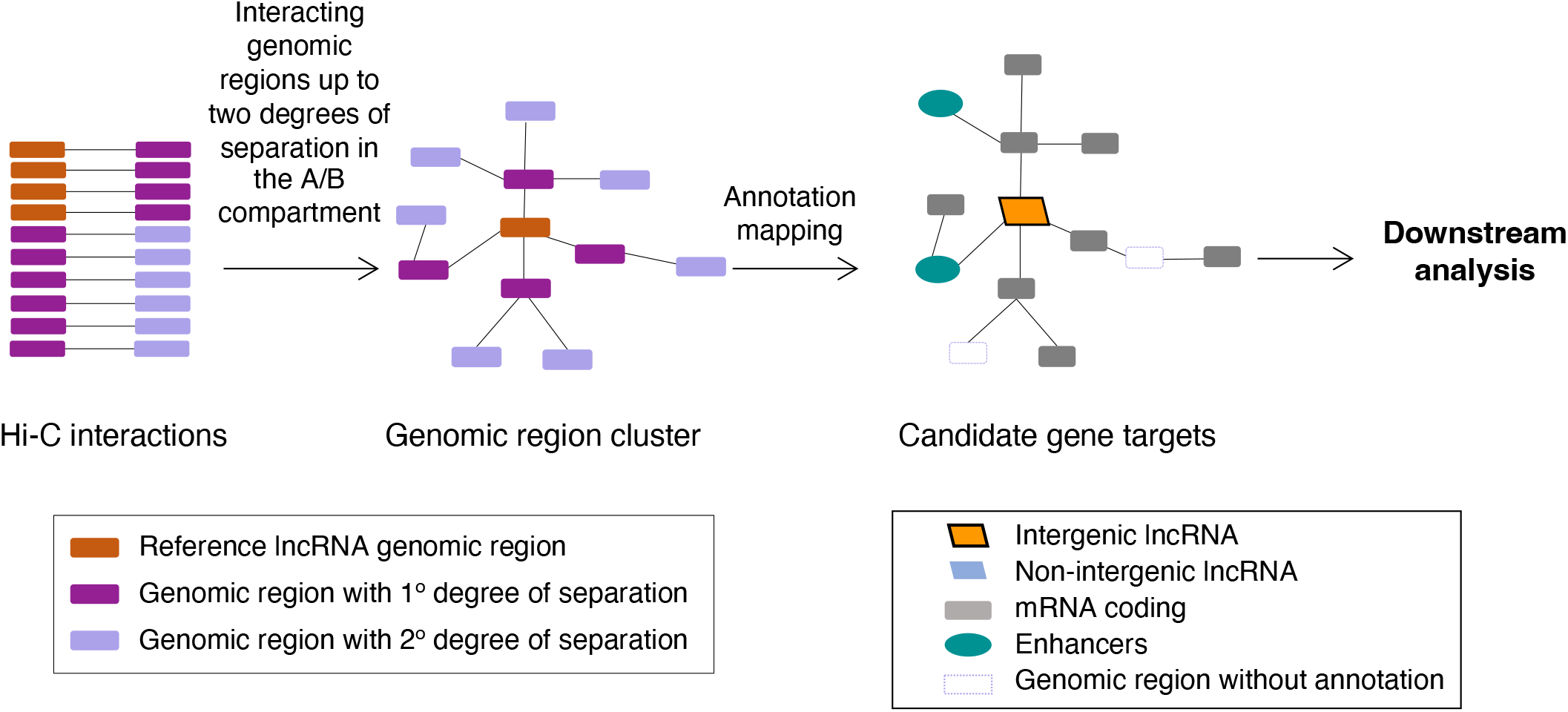
LncRNAs candidate target genes. Schematic diagram showing the workflow of lncRNA Hi-C candidate target genes identification.

Next, the correlation in expression between each lncRNA and its candidate mRNA target genes was calculated. Whereas 26.37 - 48.12% of intergenic lncRNAs and 28.59-54.88% of non-intergenic lncRNAs had a statistically significant either positive or negative average expression correlation, the majority showed no significant average expression correlation **(Figure 4)**, indicating that expression correlation analysis alone cannot reveal the clusters of associated genes found by chromatin conformation analysis. This shows that Hi-C interactions provide a unique set of candidate target genes of each lncRNA which is not completely captured by analyzing co-expression analysis.

**Figure 4:**
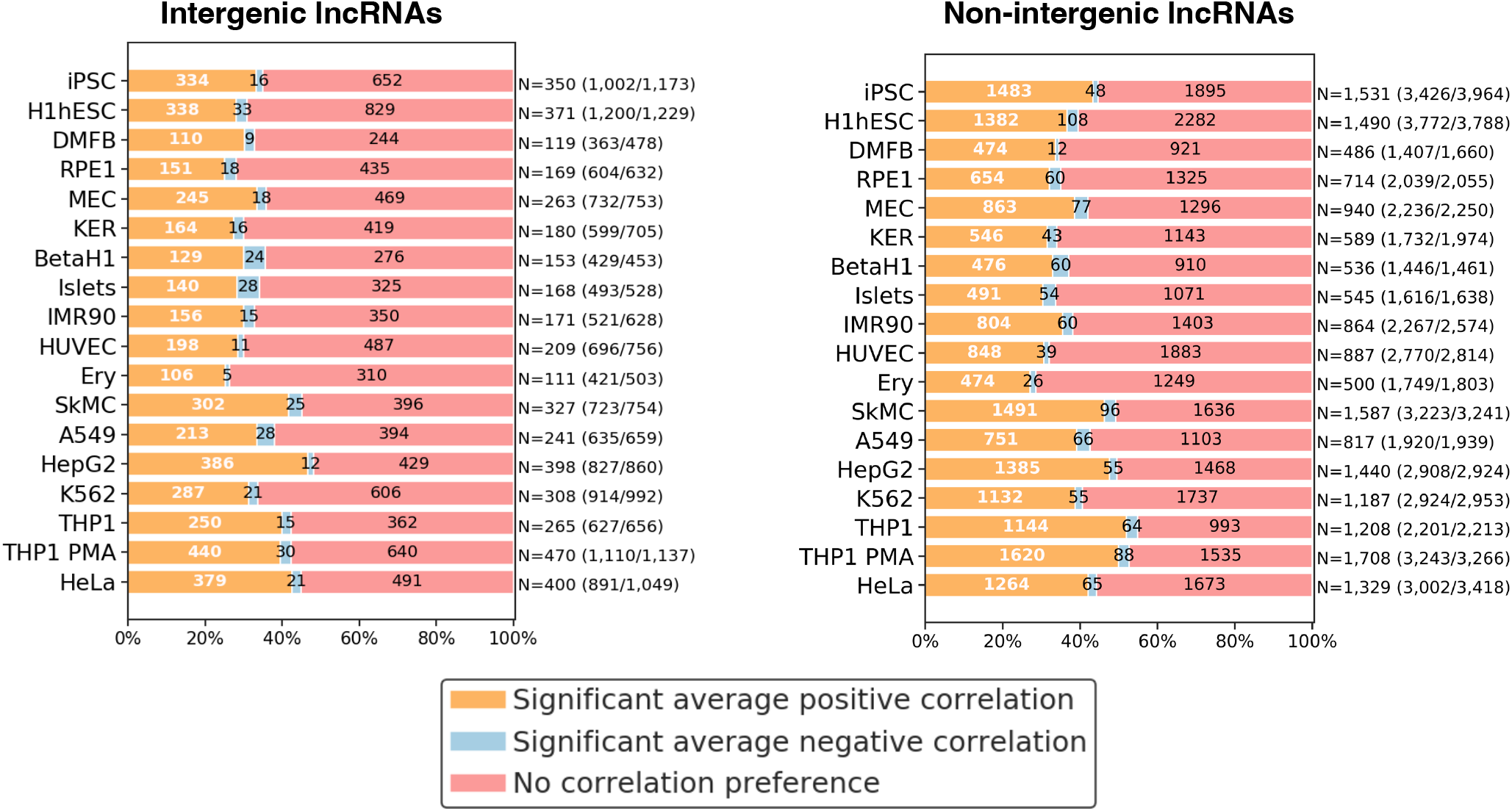
Expression correlation between LncRNAs and it’s candidate target genes. Intergenic and non-intergenic lncRNAs with significantly more candidate target genes with a positive average expression correlation, with a negative average expression correlation, or neither, in each cell type. On the right of each row, the number of intergenic lncRNAs with significant positive or negative average expression correlation with candidate mRNA target genes is shown, followed by the number of intergenic lncRNAs tested for correlation/ the number of intergenic lncRNAs expressed in that cell type in parentheses. The clusters were classified into three categories by comparing the set of expression correlation values of cluster genes to 0 using one sample Student’s t-test (P-value ≤ 0.05).

### LncRNAs as potential recruiters of RBPs at candidate target genes

There are strong evidence that lncRNAs acting in cis are known to bind to RBPs to facilitate the formation of RBP condensates to recruit the RBP to chromatin [16–18]. Based on eCLIP (enhanced CLIP) data for K562 and HepG2 cells, we found 1,754 and 1,577 lncRNA transcripts, respectively, bound by at least one RBP. The cis-score for lncRNAs to which at least one RBP bound is significantly higher than for lncRNAs without RBP indicating that lncRNA that can form RNA-chromatin interaction are likely to bind to an RBP (**Figure 5**). In K562 (cell type with RNA-chromatin, and RBP eCLIP and ChIP-seq data available), 92 reference lncRNAs have Red-C RNA-chromatin interactions with the genomic regions containing promoters with ChIP-Seq signal for at least one DNA-binding RBP. Further, eCLIP data showed that transcripts of 63 out of the 92 lncRNAs are also bound by these RBPs. Overall, this shows that lncRNAs with higher cis-score are candidate lncRNAs that are likely to facilitate recruitment of RBPs in cis to the promoter and enhancer of its candidate target genes.

**Figure 5:**
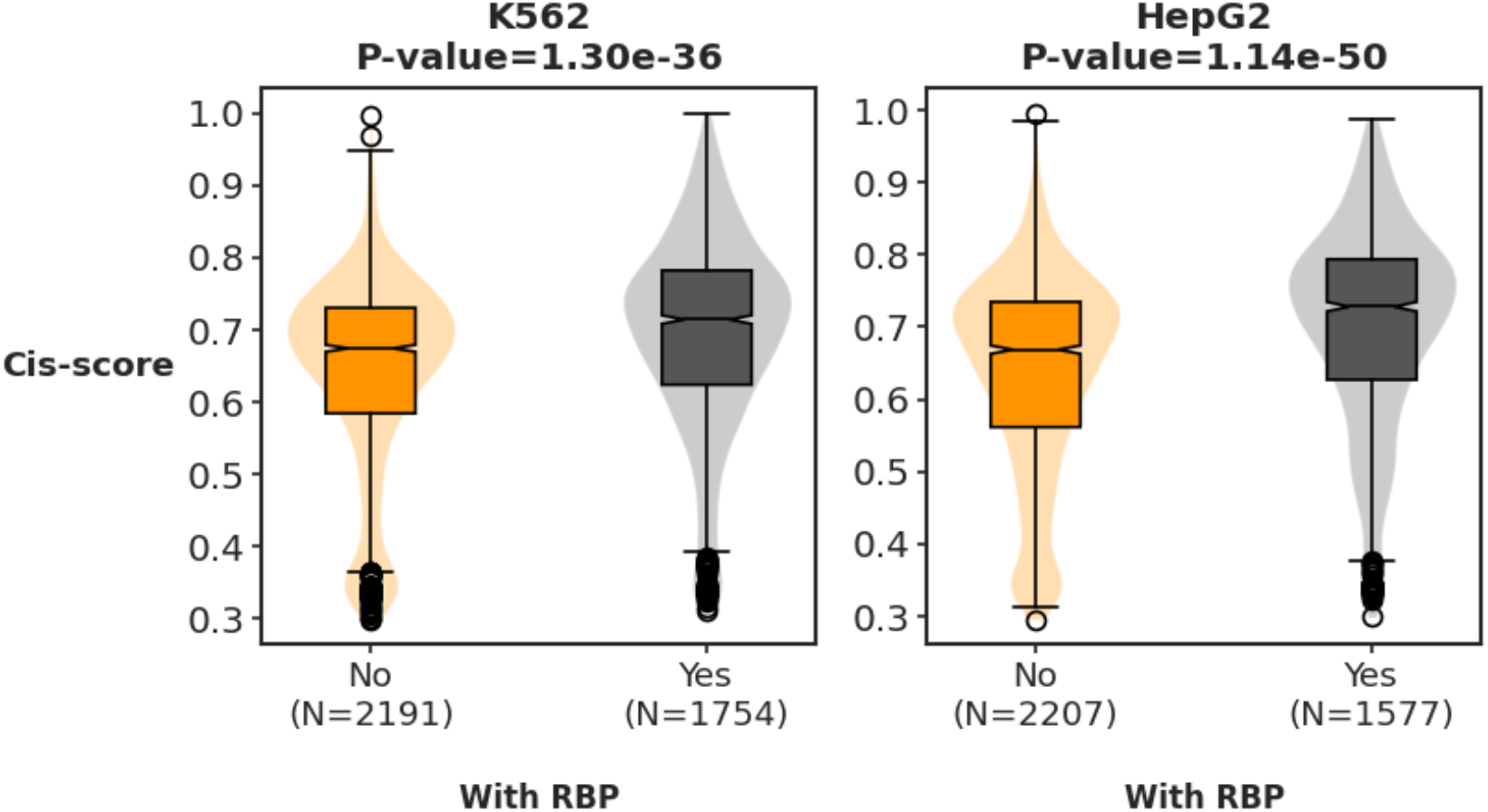
Comparison of cis-score between lncRNAs with and without RBPs binding. Each panel shows one cell type for which RBP eCLIP data is available. The significance of difference in cis-score between two classes of lncRNAs was determined using a one-tailed Mann-Whitney U test.

As an example, the currently unannotated broadly expressed lncRNA U91328.19 (ENSG00000272462) **(Supplementary Figure 6a**) with cis-score 0.52 - 0.85 has multiple histone genes as candidate target genes **(Supplementary Figure 6b; Figure 6a)**. This lncRNA has been reported to be associated with GWAS SNPs related to hay fever and eczema [47]. This is supported by the GO analysis of its candidate targets which show enrichment for terms like ‘interleukin-7-mediated signaling pathway’, ‘innate immune response in mucosa’ and ‘antibacterial humoral response’ **(Supplementary Figure 6b)**, and also GWAS heritability analysis of the A/B compartments showing enrichment for the ‘disease_ALLERGY_ECZEMA_DIAGNOSED’ trait **(Supplementary Figure 6c)**. RBPs HNRNPL, SRF1, and ILF3 which are involved in different immune response pathways [48], [49–51], are enriched at the promoters of the respective lncRNA’s candidate target genes in K562 and HepG2. eCLIP data show that RBP HNRNPL that also interacts with ILF3 binds to this lncRNA in K562. Further, RNA-chromatin data for K562 show that the lncRNA binds to the window containing the candidate target gene’s promoters **(Figure 6b)** to which these RBPs bind. The chromatin binding of lncRNA to these loci is not restricted to K562 but also seen in HUVEC, iPSC, MM1S, and HEK293T cells **(Figure 6b)** where the lncRNA is expressed, suggesting that similarly, RBPs may bind to these promoters. Overall, this indicates that the lncRNA U91328.19 may guide the recruitment of RBPs to the promoters of its candidate target genes and potentially involve in the cellular immune response.

**Figure 6:**
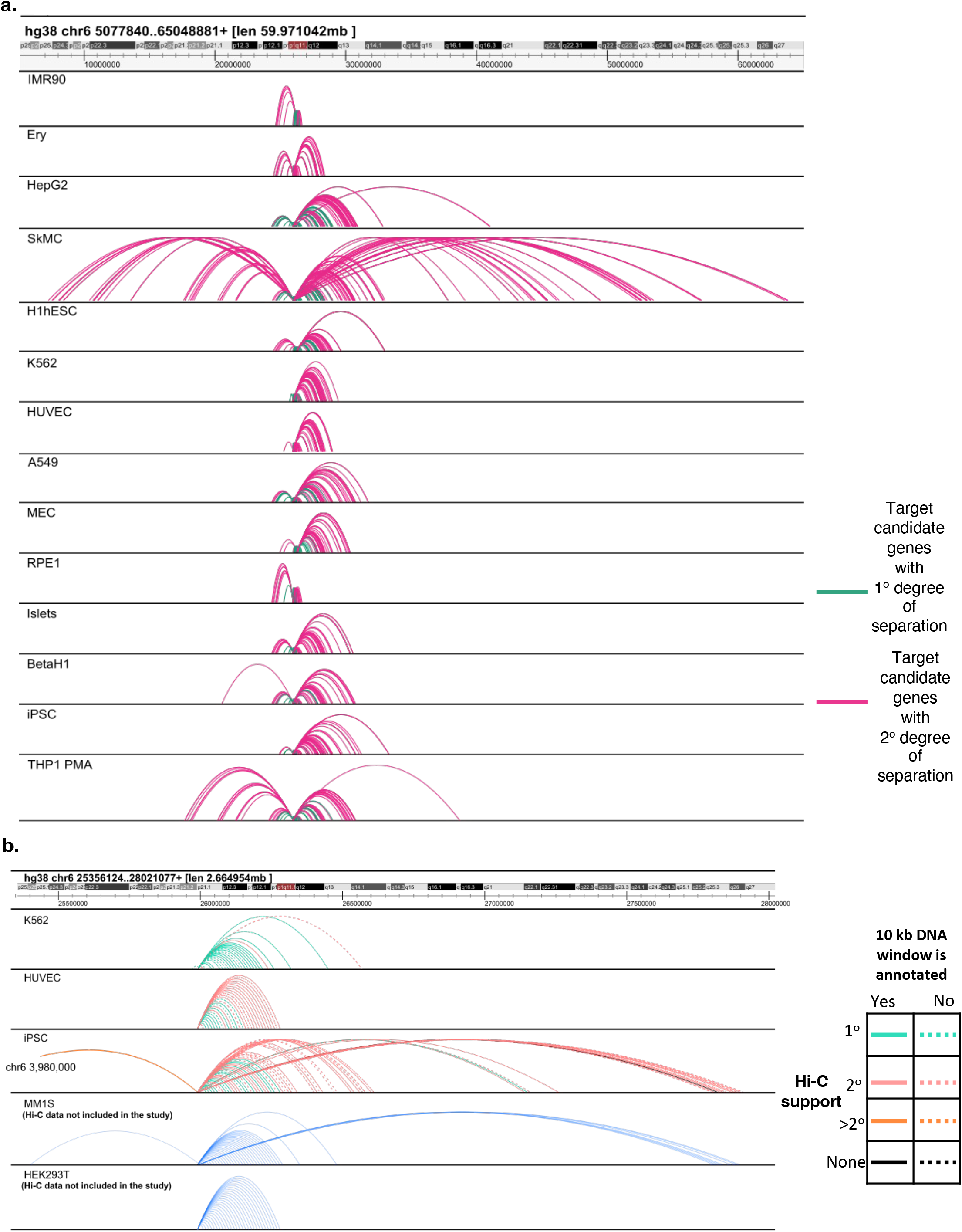
Hi-C and RNA-chromatin interactions for lncRNA ENSG00000272462. **a.** The top track shows the genomic location of the interaction followed by tracks showing the Hi-C annotated interactions between lncRNA ENSG00000272462 and its candidate target genes in different cell types. **b.** RNA-chromatin interactions for lncRNAs in different cell types. The colors of interaction tracks for K562, HUVEC, and iPSC cells show the level of support of RNA-chromatin interactions by Hi-C interaction.

### Interactive querying and visualization of lncRNA functional annotations across cell types

We created an interactive system, accessible at https://fantom.gsc.riken.jp/zenbu/reports/#F6_3D_lncRNA, to query and visualize the lncRNA properties, their candidate targets and properties derived from our analysis and to compare them across the 18 cell types. The visualization platform provides six tiers to explore the data *viz.* 1) Genome browser view, 2) Differential genomic interaction frequency table for each cell type pair, 3) Interactions associated with individual genes, 4) For each lncRNA: a. Hi-C candidate gene targets, b. associated GO terms, TFBSs, and RBPs, and c. genomic interactions are shown. and, 4) Biological features comparison among cell types for candidate targets of each lncRNA. The interactive system also allows comparing lncRNA expression, GO terms, GO genes-reference lncRNA expression correlation, genome-wide association study (GWAS) traits, and TFBS motif enrichment across the cell types.

As an example of generated data supports the previously published studies, the non-intergenic lncRNA HOTAIRM1 (ENSG00000233429) is expressed in multiple human cell types **(Supplementary Figure 7a)** with the cis-score range of 0.45-0.69. It is known to regulate the local spatial arrangement of the HOXA gene clusters [52, 53] and hence regulates the proliferation and differentiation of cells[53–55]. The candidate targets are enriched for GO terms related to differentiation (**Supplementary Figure 7b, Supplementary Figure 8a**). Similar to previously reported study[52], the expression of the lncRNA and HOXA cluster genes in this GO category are positively correlated (**Supplementary Figure 7b)** as reported in a previous study [52]. Chromosome Conformation Capture studies have shown that knocking down HOTAIRM1 results in stronger interaction with HOX1/2 and HOX9/7 genes depending on the cell type [53]. Differential genomic interaction frequency analysis between MEC and H1hESC shows that the interactions [log2(FC)=-1.17] between HOXA2 and HOXA9 are weaker in MEC (HOTAIRM1 expression: 63.47 Tag Per Million [TPM]) compared to H1hESC (HOTAIRM1 expression: 2.48

TPM) and stronger interactions [log2(FC) =1.55] when compared to SkMC (HOTAIRM1 expression: 21.03 TPM) supporting the findings of Wang *et. al.* [53]. HOTAIRM1 binds to the HOXA2 genomic window in HUVEC (HOTAIRM1 expression: 33.31 TPM), HFFc6 (HOTAIRM1 expression: 3.74 TPM; HOXA2 not expressed in HFFc6), and HEK293T (HOTAIRM1 expression: 50.21 TPM). No significant RNA-chromatin interactions were present for HOTAIRM1 around HOXA clusters in MM1S cells (HOTAIRM1 expression: 29.66 TPM) (**Supplementary Figure 8b)** suggesting the lncRNA may directly interact with chromatin in some but not all cell types. Further, TFBSs for SREBF1,2, and SPZ1 transcription factors associated with proliferation and differentiation [56–58], are enriched at the promoters of candidate targets **(Supplementary Figure 7c).** Overall, Hi-C and expression data concur with the previous findings [52], [53] regarding HOTAIRM1 demonstrating the strength of our analysis.

## Discussion

In this study, we have calauclated the cis-score as a parameter to identify the potential cis-acting lncRNAs and their putative target regions across 18 cell types. Our analysis provides a framework where using Hi-C one can identify the putative targets of a lncRNA. While several techniques have been developed in recent years that probe RNA-chromatin interactions [36–38, 59, 60], lncRNAs are underrepresented in these data due to their rapid degradation by the exosome compared to mRNAs. In contrast, Hi-C is not affected by RNA stability and is, therefore, more powerful for identifying lncRNA targets. Our Hi-C clustering strategy includes regions that are directly interacting with the lncRNA promoter as well as regions that are spatially proximal to the lncRNA but not directly interacting, and thereby takes the mobility of lncRNA transcripts in the nucleus into account. A recent microscopy study has shown that, during differentiation of mouse embryonic cells to neural progenitor cells, promoters were found to be in close proximity but not in direct contact with the enhancers regulating their activity [61]. LncRNAs can facilitate such spatial colocalization of enhancers and promoters several megabases apart by promoting chromatin reorganization, as for example demonstrated for lncRNA-CSR and Platr22 in mice [23, 24].

This systematic identification of candidate target genes is important especially for intergenic lncRNAs as their linear genomic environment lacks genes that could otherwise be used to suggest a functional role. Several lncRNAs with a biological role are known to be exosome-sensitive [23, 62]; their rapid degradation and therefore low prevalence necessitates a strategy independent of gene expression level. Our analysis does not depend on co-expression, an alternative method to predict gene function, which is strongly affected by tissue composition, technical variation, and normalization issues that negatively affect the comparability of measured gene expression levels, provides only a single annotation of lncRNAs for all cell types, and therefore cannot identify cell type-specific functions of lncRNAs [63–68]. In contrast, the Hi-C-based candidate target gene list avoids many of these pitfalls and provides target genes for each lncRNA in each cell type separately.

Our results show that many potential cis-acting lncRNAs interact with RBPs that are enriched at their target promoters. This is in agreement with a recently reported study where hundreds of lncRNA are reported to be locally enriched in mouse embryonic stem cells [20]. LncRNAs are reported to regulate the recruitments of regulatory proteins that also act as RBPs by forming condensate-like structures [16, 69, 70]. The condensate is thought to regulate the concentration of transcription factors and thereby assist the selection of promoters to be transcribed among the ones that share the same transcription factor binding sites [41, 42]. Our analysis not only provides the list of such potential lncRNAs but also provides a framework that can be employed to identify them.

Altogether, we have identified potential cis-acting lncRNAs and summarized their potential target genes and associated features in 18 human cell types into a comprehensive resource (https://fantom.gsc.riken.jp/zenbu/reports/#F6_3D_lncRNA). This includes a web application to visualize the target genes of a lncRNA in individual cell types. We anticipate that this resource will assist in narrowing the target regions of lncRNAs for further illustration of their functions.

## Conclusion

Collectively, our study provides a list of cis-acting lncRNAs and their potential targets. Our study provides a workflow that employs lncRNA sequence features together with chromatin conformation and CAGE data to identify the lncRNA potential targets. The patterns we observed by combining different data sets highlight the role of lncRNAs in cis-regulation and will assist in designing the mechanistic studies to explore the function of lncRNAs in human cells.

## Material and Methods

### RNA-chromatin interactions

We collected and reprocessed published data from the following experiments: Red-C [35], GRID-seq [38], and iMARGI [39, 40] (**Table 14**). First, we applied FastUniq [71] for filtering out possible PCR duplicates in paired-end mode. Next, we used Trimmomatic [72] to detect and remove low-quality bases in paired-end mode with default parameters except for window size (5) and base quality threshold (26). Next, we excluded the read pairs lacking the presence of experiment-specific technical sequences to consider only proper RNA-DNA chimeric reads.

Next, the RNA and DNA parts of contacts were collected in two separate files in fastq format. The RNA (longer than 13bp) and DNA (longer than 17bp) parts were independently mapped to the reference human genome (GRCh38.p13) with hisat2 [73].

The final list of RNA-DNA contacts included only the contacts with both DNA and RNA parts uniquely mapped to the reference genome with less than three mismatches, the contacts with RNA-parts mapped to splice junctions were discarded.

For all datasets, the resulting RNA parts of the contacts corresponded to the reverse complementary strand of the respective RNA genes.

### Technology-specific details

Red-C raw paired-end reads were processed as described in the original article [35]. The read pairs were pre-filtered based on the inclusion of the bridge segment in the 1st (forward) reads; for each contact, we obtained DNA, RNA-3’ (neighboring the bridge sequence), and RNA-5’. We checked that RNA-3’ and RNA-5’ parts are mapped to the opposite strands of the same chromosomes within 10 Kb from each other and considered only the RNA-3’ part in the final list of contacts. For all iMARGI datasets, the reads were pre-filtered requiring 5’ CT for the 2nd (reverse) reads; 5’ NT were allowed for HUVEC iMARGI data due to lower quality scores at 5’ ends of the reads. Additionally, the first two bases of the forward reads were removed according to the original iMARGI protocol. The GRID-seq data was already preprocessed by the authors (PCR duplicates were removed and technical bridge sequences were trimmed), thus the respective stages of our pipeline were skipped.

All genes (mRNA, intergenic, and non-intergenic lncRNA) that have at least one promoter with expression ≥ 0.5 TPM based on CAGE were used for the overlap analysis. The genome was divided into 10kb bins and the annotated RNA reads were aggregated. To reduce the effect of genomic windows with very high interaction counts (typically found near the gene where the RNA is produced), skewing the distribution, we used an iterative approach in which significant interactions are removed from the data set in each iteration. The background probability for a bin was calculated by dividing the count of trans mRNA binding in that bin by the total number of trans mRNA reads. To estimate the significance of the RNA binding in each bin, we performed a one-sided binomial test using binom_test (x,n,p) from scipy where x = the number of reads for the gene in the bin; n= total number of remaining reads for the gene, p=background probability calculated using trans-binding mRNA in the bin. The binomial test was performed iteratively until no additional interactions (FDR ≤ 0.01) were found. For each iterative step, interactions with FDR ≤ 0.01 from the previous step were removed and the number of gene reads from those bins were subtracted from n (the total number of reads for the gene). The bins where gene interaction was supported by at least 3 reads and FDR ≤ 0.01 were defined as bins where RNA from that gene binds to the chromatin significantly, and were defined as significant RNA-chromatin interactions. The genomic bins of significant RNA-chromatin interactions were annotated by mapping the strongest promoter for mRNAs, intergenic lncRNAs, and non-intergenic lncRNAs to identify RNA-chromatin gene pairs.

### Cis-score calculation

For each lncRNA, genomic bins with at least 2 RNA reads mapping to the corresponding lncRNA were used to count cis and trans bins. LncRNA with ≥75% of total bins as cis were marked as cis-binding lncRNAs. The CAGE fractionation data for cell types A549, H1hESC, HUVEC, HeLa, HepG2, IMR90, K562, KER, MCF7 (MEC), SK-N-SH, and iPSC was used to calculate the expression of each in gene in the nuclear and cytoplasmic fractions (TPM). The formula was used to define the fraction of lncRNA in the nucleus compared to the cytoplasm log2((nuclear + cytoplasmic expression)). For the cell types where fractionation data was not available, the mean fractionation values were assigned. All the exons from the transcripts associated with the expressed promoters were selected to search for the sequence features. For the SIRLOIN motif representative motif [CT][GA]CCTCCC[GA][GA]GTTCAAG[CT]GAT[TC]CTCCT[GA]CCTCAGCCTCCCGA, derived from [32] and for the U1 motif representative sequence, CAGGTGAGT was searched in the selected exons using function fuzznuc of EMBOSS package. The repeatmasker track from UCSC was used to determine repeat frequency and invert Alu repeats. The cis-score (probability of lncRNA to be cis) was predicted by using binary logistic regression and using nuclear fraction, log value of expression level, frequency of SIRLOIN motif, U1 motif, repeat sequences, and inverted Alus as parameters.

### Hi-C data generation and processing

iPSC Hi-C data was generated as described in Ramilowski *et al.* [74] while data for the remaining cell types were obtained from previously published studies (**Table 5**). Data for each replicate was processed using HICUP ver. 0.5.10 [75] which involved read truncation, mapping, filtering experimental artifacts, and de-duplication. The alignment files for all replicates for each cell type were merged to perform downstream analysis. GRCh38 primary human genome assembly (hg38) was used for the analysis.

### Gene and enhancer models and primary annotations

FANTOM CAT gene models [1] and hg38 FANTOM 5 bidirectional enhancers [76] (FANTOM-5 enhancers) (https://zenodo.org/record/556775) were used as the primary genome annotation. The FANTOM CAT gene classes used in this study are mRNAs (protein-coding), intergenic lncRNAs, antisense lncRNAs, divergent lncRNAs, and sense intronic lncRNAs, with the latter three classes collectively referred to as non-intergenic lncRNAs.

### The expression level of promoters and bi-directional enhancers

Expression levels of promoters and enhancers were determined using CAGE data. iPSC CAGE data was generated for this study using the nAnT-iCAGE protocol [34], while publicly available matched CAGE libraries were used for the remaining cell types **(Table 15)**. Promoter CAGE tag counts were estimated by intersecting CTSS files for individual libraries with the promoter bed file using bedtools (ver. 2.26.0) and were normalized to calculate promoter expression in tags per million (tpm). The expression for a promoter in a cell type was determined by calculating the mean expression across all CAGE libraries for that cell type. For each gene, the promoter with the highest expression level, requiring a minimum expression of 0.5 tpm, and minimum 3 tag counts in at least one CAGE library was used for the downstream analysis. Next, the CAGE expression of enhancers was calculated by summing the CAGE tag counts across the libraries for each cell type, ignoring the tag directionality. All enhancers with an aggregate tag count of at least 5 were used for the downstream analysis.

In the absence of matched CAGE libraries, FANTOM5 data for pancreatic tissues was repurposed for Islets and BetaH1 cells. First, expressed transcripts in Islets and BetaH1 cells were determined using RNA-seq data from published studies [77, 78] (**Table 15**) using Kallisto ver. 0.45.0 [79]. Next, the strongest promoter for each gene was determined based on expressed transcripts in each cell type and pancreatic tissue CAGE data. Expressed enhancers in pancreatic tissues were assigned to both Islets and BetaH1 cell types.

### Promoter types

The precalculated chromatin state ChromHMM models for the selected cell types were downloaded from the previous studies listed in **Table 15**. The strongest promoters were intersected with corresponding cell type chromatin state models and were assigned a promoter type: H3K4me3 enriched (canonical promoters), H3K4me1 enriched (enhancer-like promoters), or Neither (undetermined) depending on the overlapping state **(Table 15)**. As chromatin state data were not available for THP1, THP1-PMA, and RPE-1, promoter types provided by FANTOM-CAT [1] were used to annotate promoters in these cell types **(Table 15)**.

### A/B compartments, TADs, and loops

Hi-C alignment .bam files for individual replicates and merged data were converted into .hic format using an in-house awk script and the Pre command from the Juicer package [80]. A/B compartments were identified at 1 Mbp resolution using the function eigenvector from Juicer package with options: -p VC <HIC file> <CHROMOSOME name> BP 1000000. The positive value represents compartment A (transcriptionally active compartment) and the negative value represents compartment B (a compartment with lower transcriptional activity compared to compartment-A). The compartments were redefined by reassigning the signs (+/-) to eigenvalues in cases where average expression values (determined using CAGE peaks) of compartment B (negative value) was higher than compartment A (positive value). Further, TADs and chromatin loops were calculated as a resource for the research community using functions from the Juicer package.

### Determining significant genomic interactions

The intra-chromosomal genomic interactions were identified using the Bioconductor package GOTHiC [81]. The Hi-C data across biological replicates for each cell type was merged and statistically significant cis-genomic interactions were identified at 10 kbp resolution. The alignment .bam files were converted into .gothic files using the format conversion script hicup2gothic from the HiCUP package. The interactions were calculated using .gothic together with corresponding restriction enzyme files **(Table 5)** for each cell type. All the interactions supported by at least 5 read pairs and q-value ≤ 0.05 were defined as significant genomic interactions. The number of genomic interactions per cell type varied from 2,540,361 to 46,975,256 with Hi-C sequencing depth (**Table 7**) and included interactions in both A and B compartments (compartment-A are genomic regions with higher transcription activity compared to compartment-B genomic regions) (**Table 7**). The interactions were annotated by overlaying the expressed promoters and enhancers in the selected cell types to identify the promoters interacting with each other. In cases where promoters for more than one gene overlapped the same 10 kbp region, the interactions were counted multiple times, with one interaction for each gene. Interactions with annotations on both sides were used for the downstream pairwise analysis. The number of annotated cis-interactions varied from 50,170 to 604,677 **(Table 7)** among the cell types.

### Differential Hi-C interactions analysis

An interaction read count table (10 kb resolution) for individual Hi-C replicates was generated using straw (ver. 0.0.8). Islets and BetaH1 data were excluded from the analysis as they had only one Hi-C library. The pairwise differential Hi-C analysis was performed using the Bioconductor package multiHiCcompare (ver. 1.8.0). All genomic interactions with 5 read counts in at least two Hi-C libraries were tested for the differential interactions. Interactions with **|**log2(Fold change)**|** ≥ 1 and FDR-corrected P-value ≤ 0.1 were defined as differential Hi-C interactions. Enrichment of H3K4me1 promoters with differential interactions compared to H3K4me3 + Neither promoters was tested using one-tailed Fisher’s exact test for individual gene classes separately. The promoters that had different promoter types between two cell types were excluded from the analysis. All promoters that overlapped with interactions tested for differential Hi-C were used for the analysis. The relation between Hi-C interaction, mRNA, and intergenic lncRNA expression change was determined using the Chi-squared test. For H3K4me1 and H3K4me3 enriched promoters of each gene class and enhancers, Spearman correlation was calculated between change in the differential Hi-C interactions and log2 (fold change) in the expression of interacting mRNA gene in query cell type compared to reference cell type. Enrichment of transcription factors in differential Hi-C interactions was performed for each cell-type pair using one-tailed Fisher’s exact test. The list of human transcription factors by Ravasi *et al.* [82] was used for the analysis.

### Hi-C clusters

Hi-C genomic clusters for each lncRNA were defined using significant genomic interactions. The genomic window overlapping with the reference lncRNA promoter was selected as the reference genomic region. To define the Hi-C genomic clusters, all the genomic regions connected to the reference genomic region by Hi-C interaction up to 2 degrees separation [reference lncRNA genomic region → genomic region that have Hi-C interaction connecting to the reference lncRNA genomic region (Genomic region with 1° degree of separation from reference genomic region) → genomic regions that are connected to Genomic region with 1° degree of separation by Hi-C interactions (Genomic region with 2° degree of separation from reference window)] within the A or B compartment (extended by +/-100 kbp) in which the reference lncRNA regions is situated were used to define the Hi-C genomic region clusters **(Figure 3)**. In the case of DMFB, Ery, and HUVEC where the Hi-C sequencing depth was relatively low, we also included genomic windows connected within 3 degrees of separation. The expressed promoters and enhancers were mapped to the Hi-C genomic region clusters to identify the potential target genes **(Figure 3).**

### Gene ontology (GO) enrichment analysis

The GO term database from NCBI was downloaded on Nov 28^th^, 2019 (file: gene2go from ftp://ftp.ncbi.nlm.nih.gov/gene/DATA/). All mRNA genes with an entrezID that are expressed in a given cell type and belong to at least one Hi-C cluster were used for the analysis. The GO term enrichment analysis for each Hi-C cluster was performed using a one-sided Fisher’s exact test (details are given in **Table 16**). The background consisted of all genes in all clusters other than the genes in the reference lncRNA cluster. The analysis was performed for the GO terms that have at least one mRNA gene in their geneset in common with the cluster. The P-value was corrected for multiple testing using the Benjamini–Hochberg false discovery rate (FDR) multiple testing correction method. All GO terms with FDR-adjusted P-value ≤ 0.1 and at least 3 mRNA genes in their geneset in common with the cluster were defined as significant GO terms. GO term enrichment analysis was performed for the linear clusters for the “biological process” GO term category in the same way as for the Hi-C gene clusters. Further, GO enrichment analysis was also performed for gene sets provided by the Broad Institute [83–85] as a resource for annotating the lncRNA Hi-C gene clusters. Analysis was performed for all the Hi-C gene clusters with at least three mRNA genes with an entrezID. Similar to NCBI GO gene sets, enrichment analysis was also performed using EnrichR gene sets (downloaded on March 4^th^, 2021 from https://maayanlab.cloud/Enrichr/#stats) for Hi-C gene clusters.

The gene family member list from HGNC was downloaded on February 10, 2021 (https://.genenames.org/). Gene family members were mapped to GO genes based on their Ensemble geneID and for every lncRNA, the number of genes in each GO term that overlap with each gene family was counted. LncRNAs with at least one GO term that have three or more GO genes overlapping with a single gene family were classified as associated with a gene family.

### Hi-C gene cluster expression correlation analysis

For each cluster, the Spearman correlation was calculated for gene expression between the reference lncRNA and mRNA genes in the cluster across the 18 cell types. To determine if the clusters have a preference for genes that have positive or negative expression correlation with the reference lncRNA, a one-sample Student’s t-test was performed. The Hi-C gene cluster with P-value ≤ 0.05, and t-statistic value positive or negative were categorized as clusters with positive expression correlation or negative expression correlation respectively, or otherwise with no preference. The analysis was performed in the same manner including only GO-annotated mRNA genes.

### TFBSs enrichment analysis and motif correlation

Genome-wide TFBS predictions for SwissRegulon motifs [86–88] were downloaded for the hg38 human genome assembly. For each cell type, the predicted TFBSs were intersected with promoter and enhancer regions extended by +/-250 bp. In cases where multiple TFBSs for the same motif overlapped with a promoter or enhancer, the posterior probability scores of the predicted TFBSs were summed. The TFBSs with aggregate scores of at least 0.1 were assigned to the promoters and enhancers. Motifs with predicted TFBSs overlapping with less than 200 promoters and enhancers were excluded from the analysis. The significance of motif enrichment in the Hi-C gene cluster was calculated using a one-sided Fisher’s exact test. The contingency table used for analysis is described in **Table 17**. The P-values were corrected for multiple testing using the Benjamini–Hochberg multiple testing correction method. All clusters where the number of promoters + enhancers was at least 5 were tested for enrichment. All motifs with FDR adjusted P-value ≤ 0.1 and at least 3 promoters and enhancers in the cluster with a TFBS for the motif were defined as significantly enriched.

### GC content analysis

For each cell type, the GC content of promoters extended by +/-250 bp was determined using the function GC from the Bio.SeqUtils module in Biopython (ver. 1.76) [89]. The Spearman correlation between the GC content of each promoter and the number of annotated interactions was calculated for every gene class in each cell type. For the cluster analysis, the median GC content for each cluster with 5 or more promoters was calculated. To calculate the Z-score for each cluster, we randomly selected 100 sets of promoters from the set of all expressed promoters in the cell type, with the same number of promoters as in the cluster, and calculated the median GC content for each randomized set. The mean and standard deviation of these 100 GC content values was used to calculate the Z-score for the GC content of the cluster.

### RBP enrichment analysis

*Overview and analysis of ENCODE eCLIP data:* ENCODE eCLIP data for cell lines HepG2 and K562 was used for the analysis. In total, there were 234 <RBP, cell type> pairs. The data were analyzed as follows: (1) reads were preprocessed as in the original eCLIP pipeline [90], (2) trimmed reads were mapped to the hg38 genome assembly with F6 CAT genome annotation using *hisat2* [73], (3) the aligned reads were deduplicated [90], (4) the uniquely mapped and correctly paired reads were filtered with *samtools* [91], (5) gene-level read counts in exons were obtained with *plastid* [92], (6) differential expression analysis against matched controls was performed using *edgeR* [93]. Based on (6) reliable RNA targets of each RBP were defined as those passing 5% FDR and log_2_FC > 0.5. RBPs with fewer than 15 targets in Hi-C clusters were excluded from the downstream analysis resulting in the final set of 207 <RBP, cell type> pairs.

### Overview and analysis of ENCODE ChIP-Seq data

Optimally thresholded ChIP-Seq peaks of 18 and 26 RBPs for HepG2 and K562, respectively were downloaded from ENCODE and merged across replicates. U2AF1 (in K562) was excluded from analysis as having less than 15 RNA targets in eCLIP data. To annotate promoters and enhancers with ChIP-Seq peaks we required non-zero overlap between the respective genomic regions and peaks.

### Analysis of Hi-C gene cluster enrichment with RBP targets

Right-tailed Fisher’s exact test (on promoters and FANTOM5 enhancers) was used to identify Hi-C gene clusters enriched with RBP targets. Benjamini-Hochberg (FDR) correction for multiple tested RBPs was applied and cases passing FDR adjusted P-value ≤ 0.1 and with at least 3 RBP targets were considered statistically significant.

### Heritability enrichment analysis

We used stratified linkage disequilibrium (LD) score regression (ldsc software ver. 1.0.0, https://github.com/bulik/ldsc) [94] to partition the common (minor allele frequency (MAF) > 5% in European 1000 Genomes Project Phase 3 data) SNP heritability for 47 UK Biobank traits and diseases (https://data.broadinstitute.org/alkesgroup/UKBB) and four diseases which were used by O’Connor *et al.* [95–99]. We tested partitioned heritability of each Hi-C annotation (converted to hg19 genome build using UCSC liftOver tool) conditioning on the baselineLD model ver. 2.2 (https://data.broadinstitute.org/alkesgroup/LDSCORE/1000G_Phase3_baselineLD_v2.2_ldscore s.tgz). We calculated the significance of the regression coefficient for the Hi-C annotation using the Z-score.

---

**Supplementary Figure 1:**
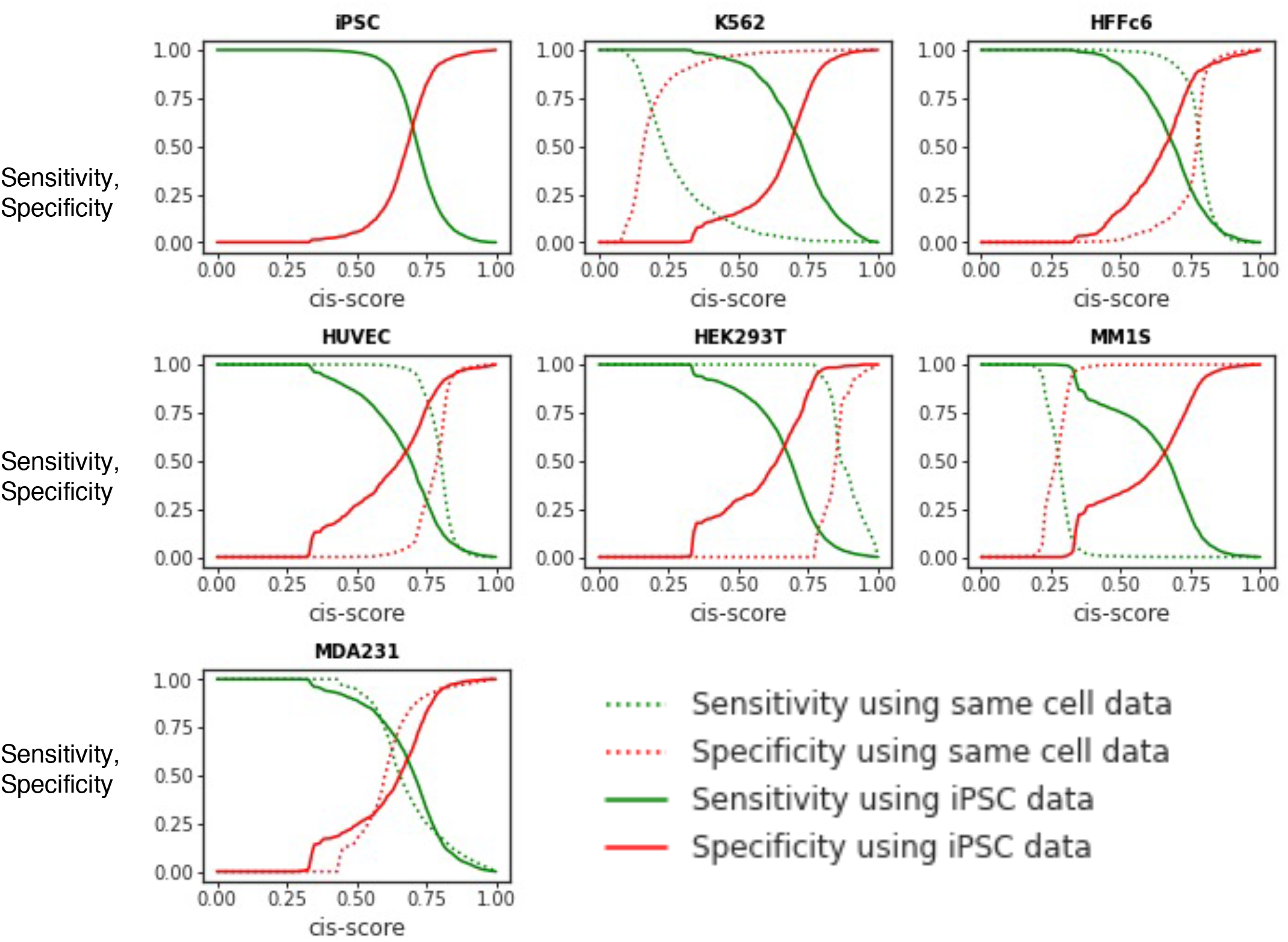
Comparison of sensitivity and specificity of cis-score predicted by model generated using data from same cell type and iPSC. Each panel shows the cell type for which RNA-chromatin data is available.

**Supplementary Figure 2:**
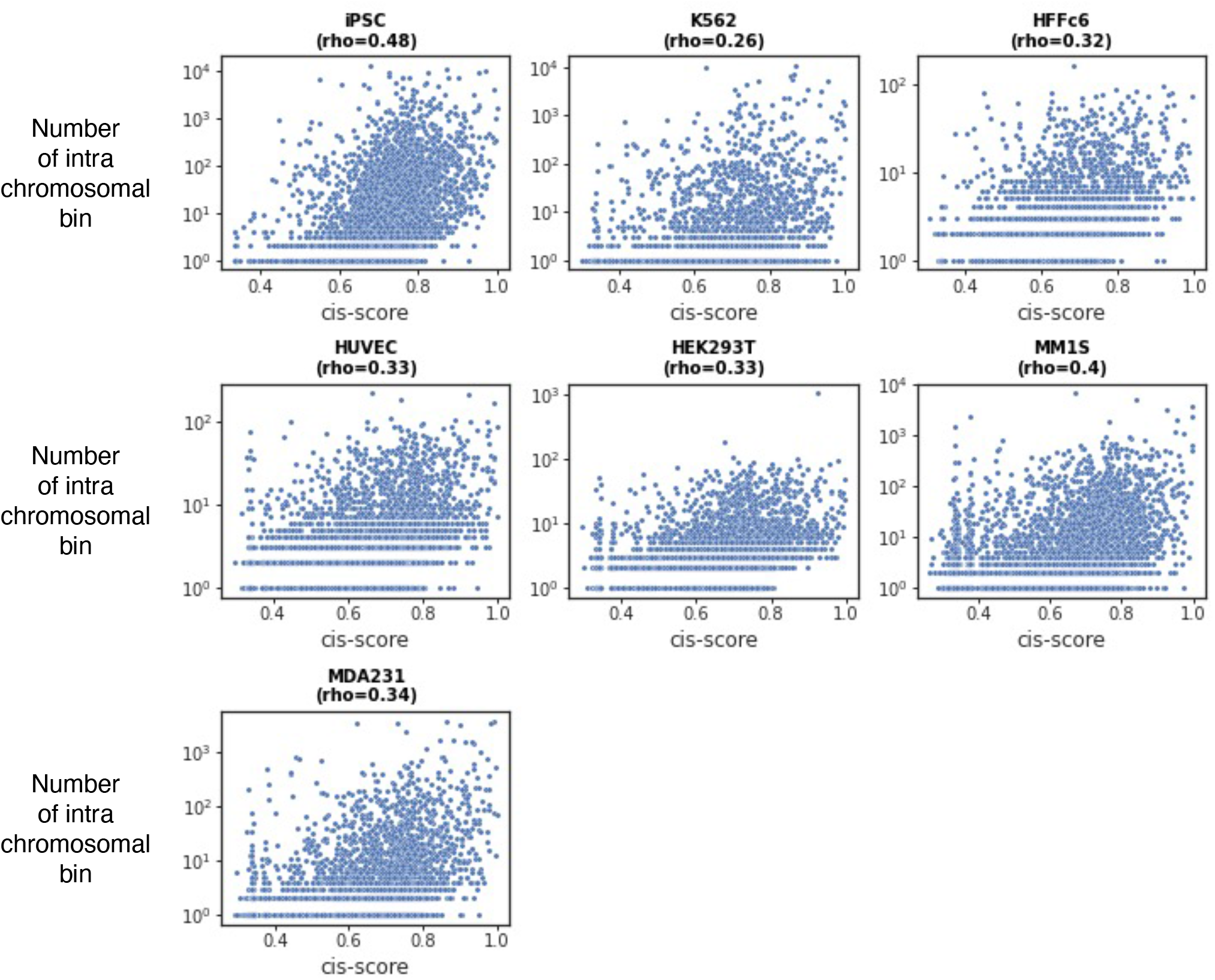
Correlation between cis-score and number of genomic bins with raw RNA-chromatin interactions. The horizontal and vertical axis shows the cis-score and number of 10kb genomic bins with cis-RNA-chromatin interaction respectively. Each panel shows one cell type, with a dot representing one lncRNA. The spearmann correlation (rho) between the cis-score and the number RNA-chromatin interaction bin is shown in the title.

**Supplementary Figure 3:**
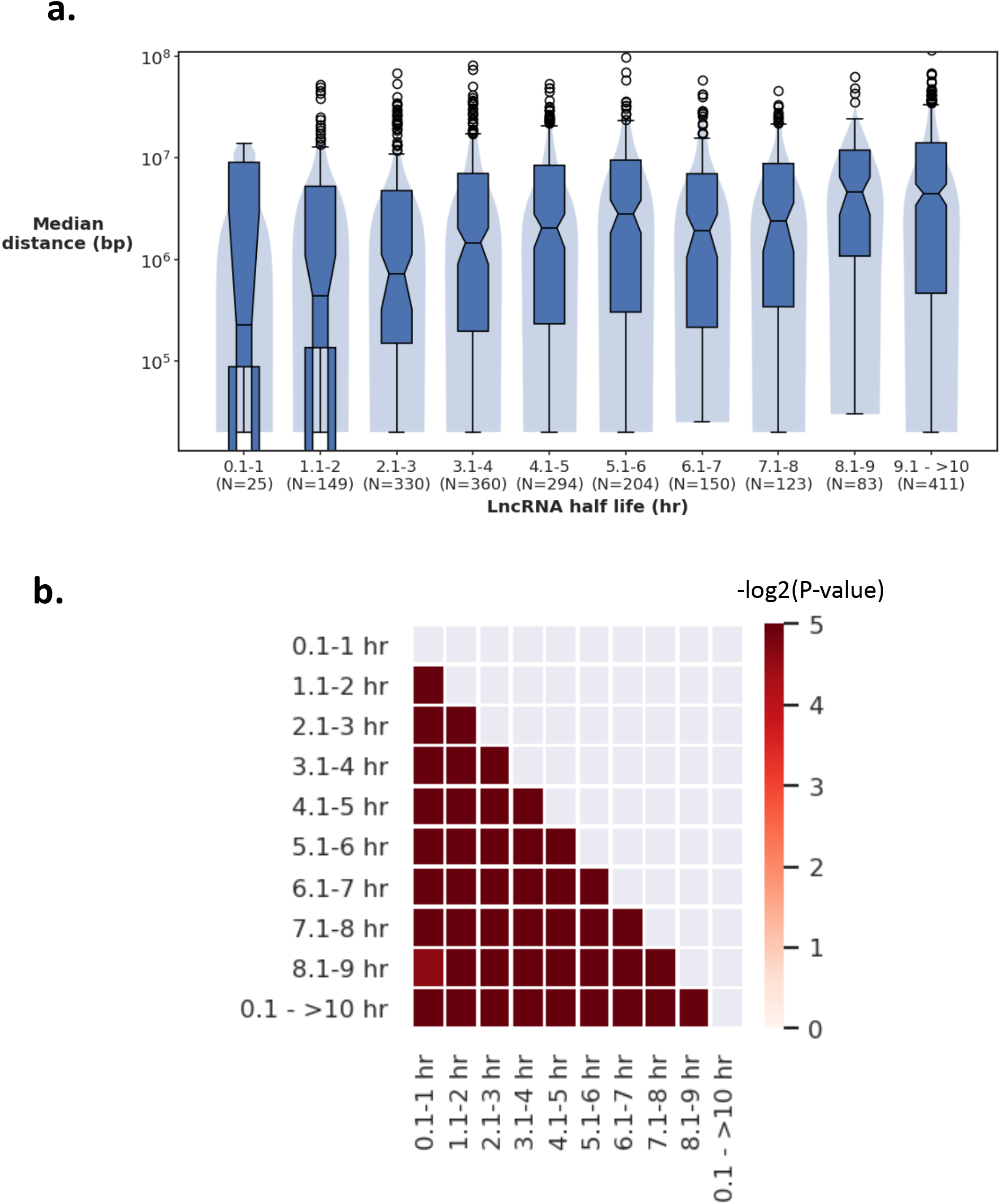
Half life of lncRNA vs median linear distance between RNA-chromatin interaction window and RNA promoter containing window. **a)** The comparison of median linear distance among different ranges of lncRNA half-life. **b)** Significance of difference of median distance values between two half-life ranges of lncRNAs was determined using the two-tailed Mann-Whitney U test.

**Supplementary Figure 4:**
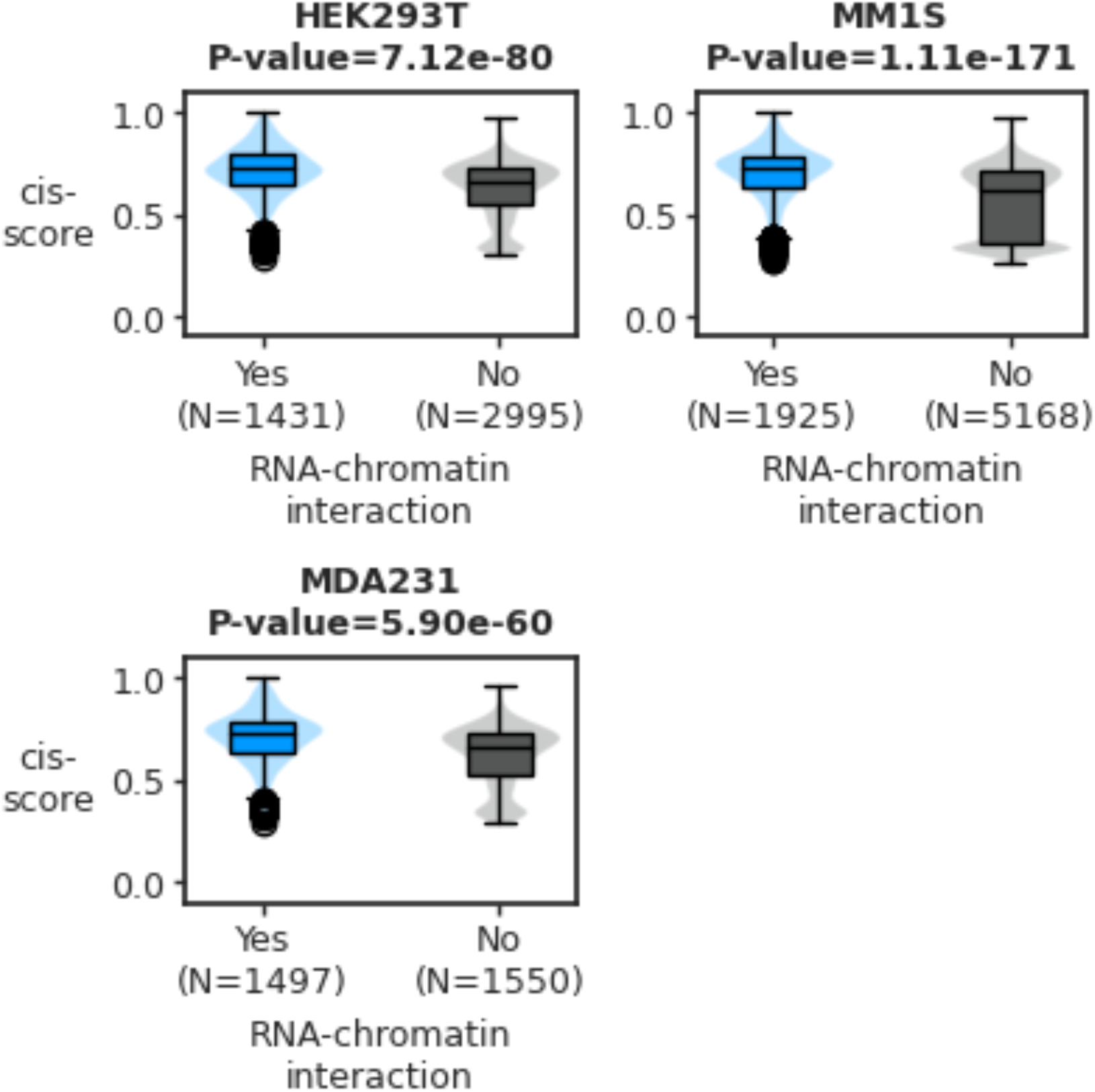
Comparison of cis-score between lncRNAs with and without atleast one RNA-chromatin interaction. Each panel shows one cell type for which RNA-chromatin data is available. The significance of difference in cis-score between two classes of lncRNAs was determined using a one-tailed Mann-Whitney U test.

**Supplementary Figure 5:**
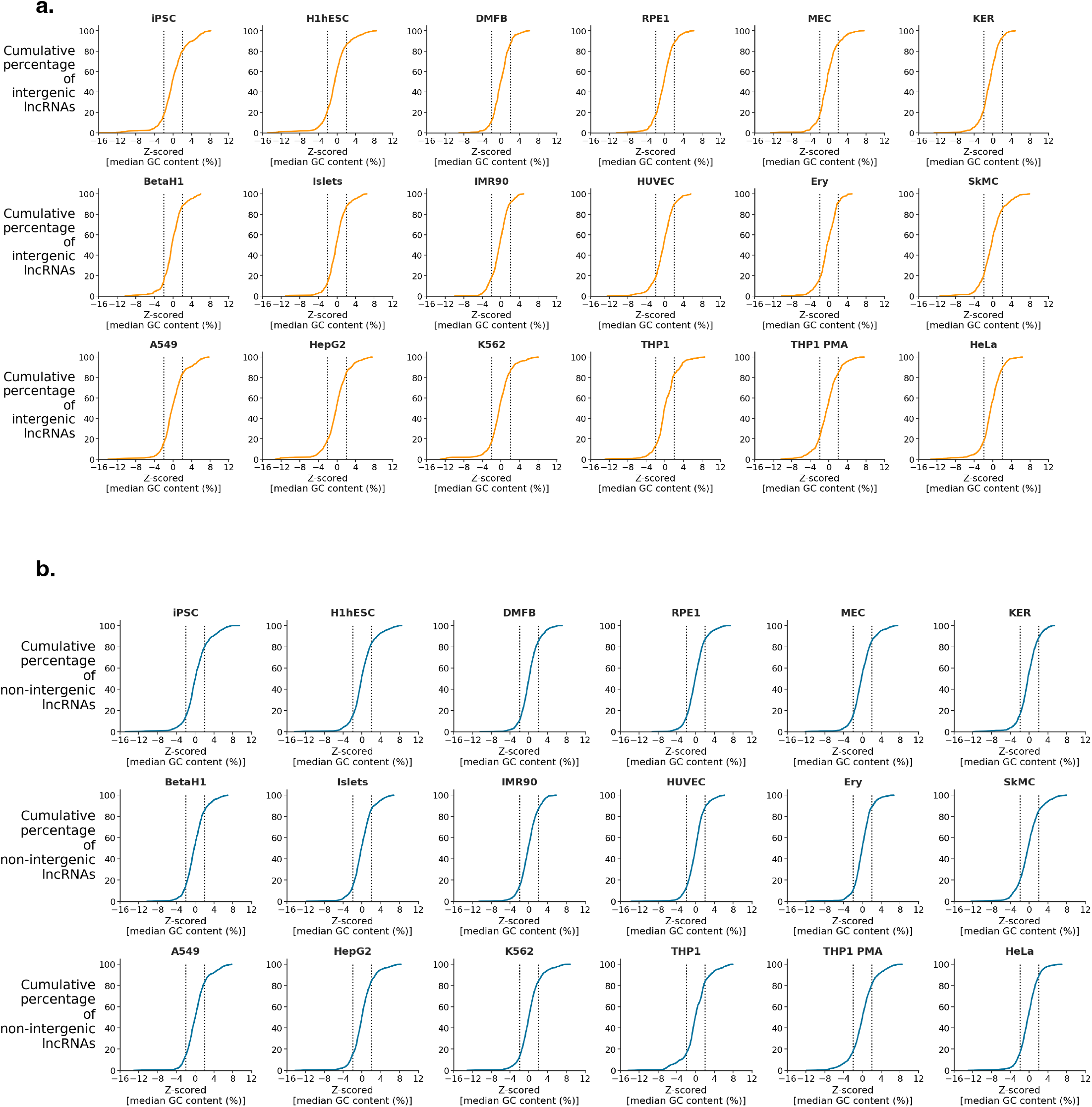
Variation in the average GC content of the HiC candidate target genes of a. intergenic lncRNAs and b. non-intergenic lncRNAs. Each plot shows the cumulative distribution for Z-scored median GC content of all lncRNAs candidate target genes in a cell type. Dashed lines indicate Z = −2 and Z = +2, corresponding to p = 0.023 and p = 0.97, respectively, for a normal distribution.

**Supplementary Figure 6:**
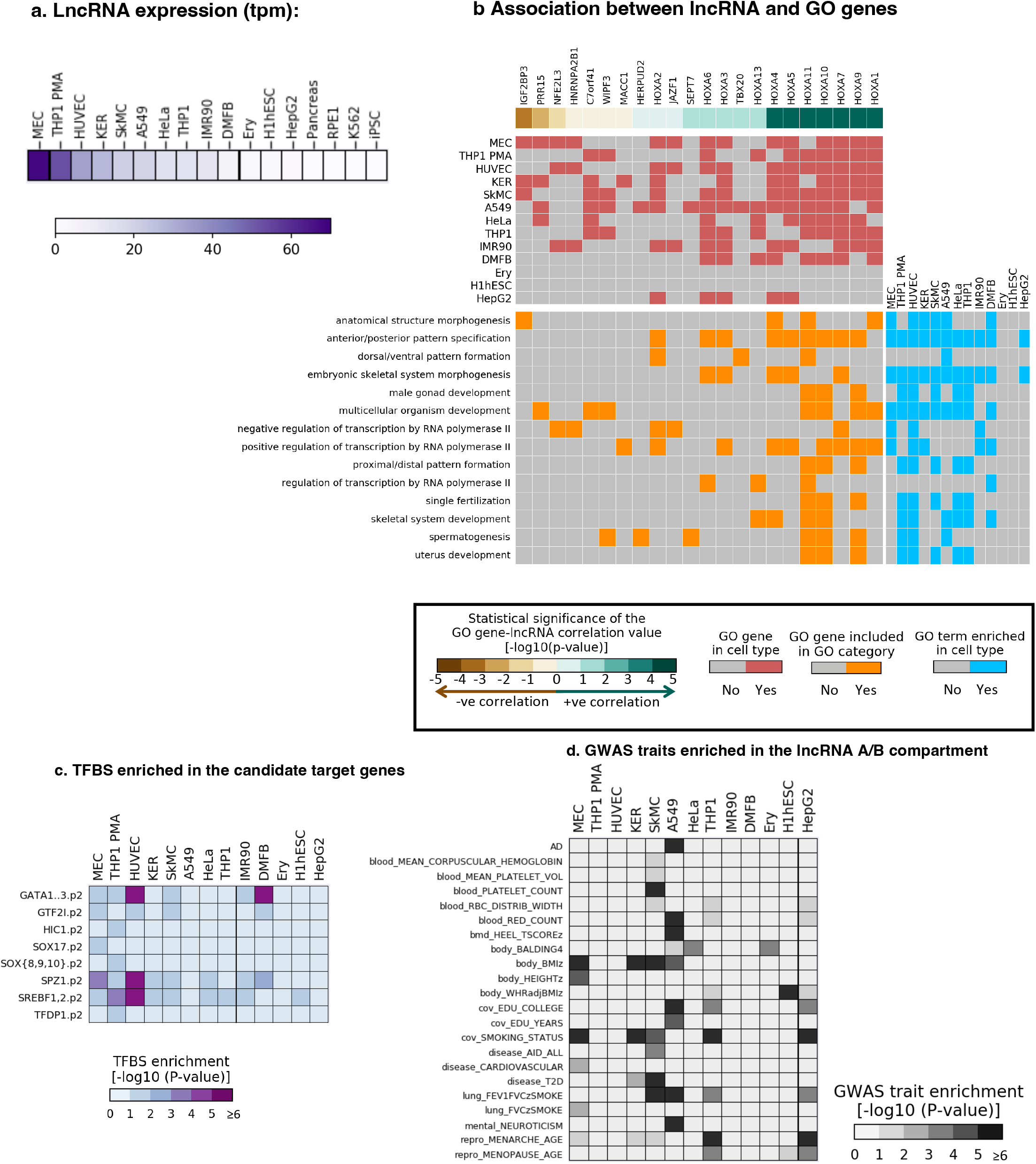
Annotation results for lncRNA ENSG00000233429. **a.** Expression of lncRNA in all 17 cell types (CAGE data from pancreas were used for both BetaH1 and islet cells). **b.** GO annotation results. The topmost heatmap shows the expression correlation between the lncRNA and GO-annotated genes in the candidate target genes. For readability, the gene names are shown on the top of the heatmap in the same order as in the heatmap. The middle heatmap shows whether the candidate target genes in each cell contains each of the GO-annotated genes. The bottom heatmap shows the membership of each GO gene in each GO category, and the heatmap on the right shows whether each GO term is enriched in each cell type. **c.** Motifs enriched (FDR adjusted P-value ≤ 0.1) in the lncRNA’s candidate target genes. **d.** GWAS trait enriched (FDR adjusted P-value ≤ 0.1) in the A/B compartment overlaps with the lncRNA’s candidate target genes.

**Supplementary Figure 7:**
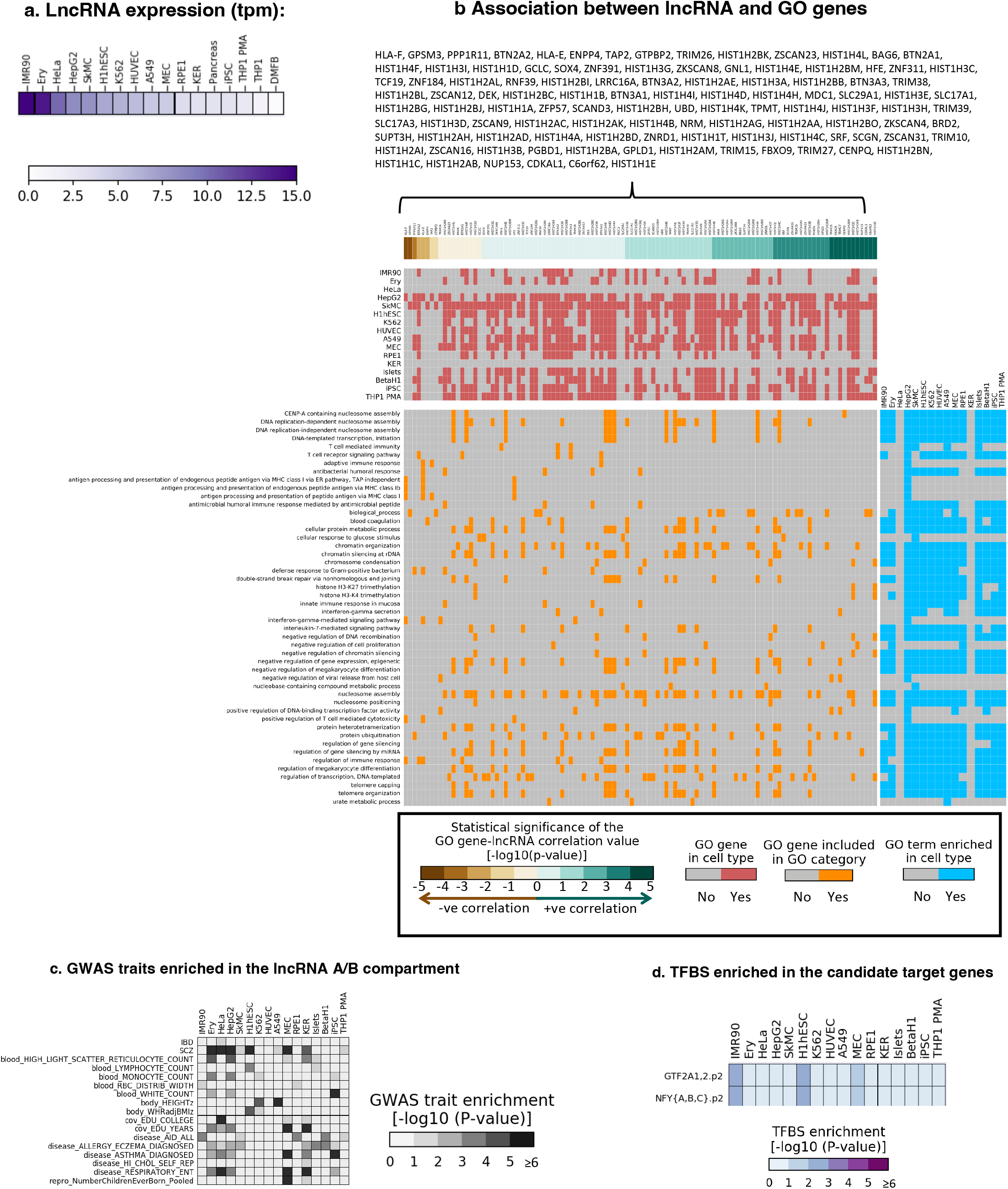
Annotation results for lncRNA ENSG00000272462. **a.** Expression of lncRNA in all17 cell types (CAGE data from pancreas were used for both BetaH1 and islet cells). **b.** GO annotation results. The topmost heatmap shows the expression correlation between the lncRNA and GO-annotated genes in the candidate target genes. For readability, the gene names are shown on the top of the heatmap in the same order as in the heatmap. The middle heatmap shows whether the candidate target genes in each cell contains each of the GO-annotated genes. The bottom heatmap shows the membership of each GO gene in each GO category, and the heatmap on the right shows whether each GO term is enriched in each cell type. **c.** GWAS trait enriched (FOR adjusted P-value,;; 0.1) in the A/B compartment overlaps with the lncRNA’s candidate target genes. **d.** Motifs enriched (FD R adjusted P-value,;; 0.1) in the lncRNA’s candidate target genes.

**Supplementary Figure 8:**
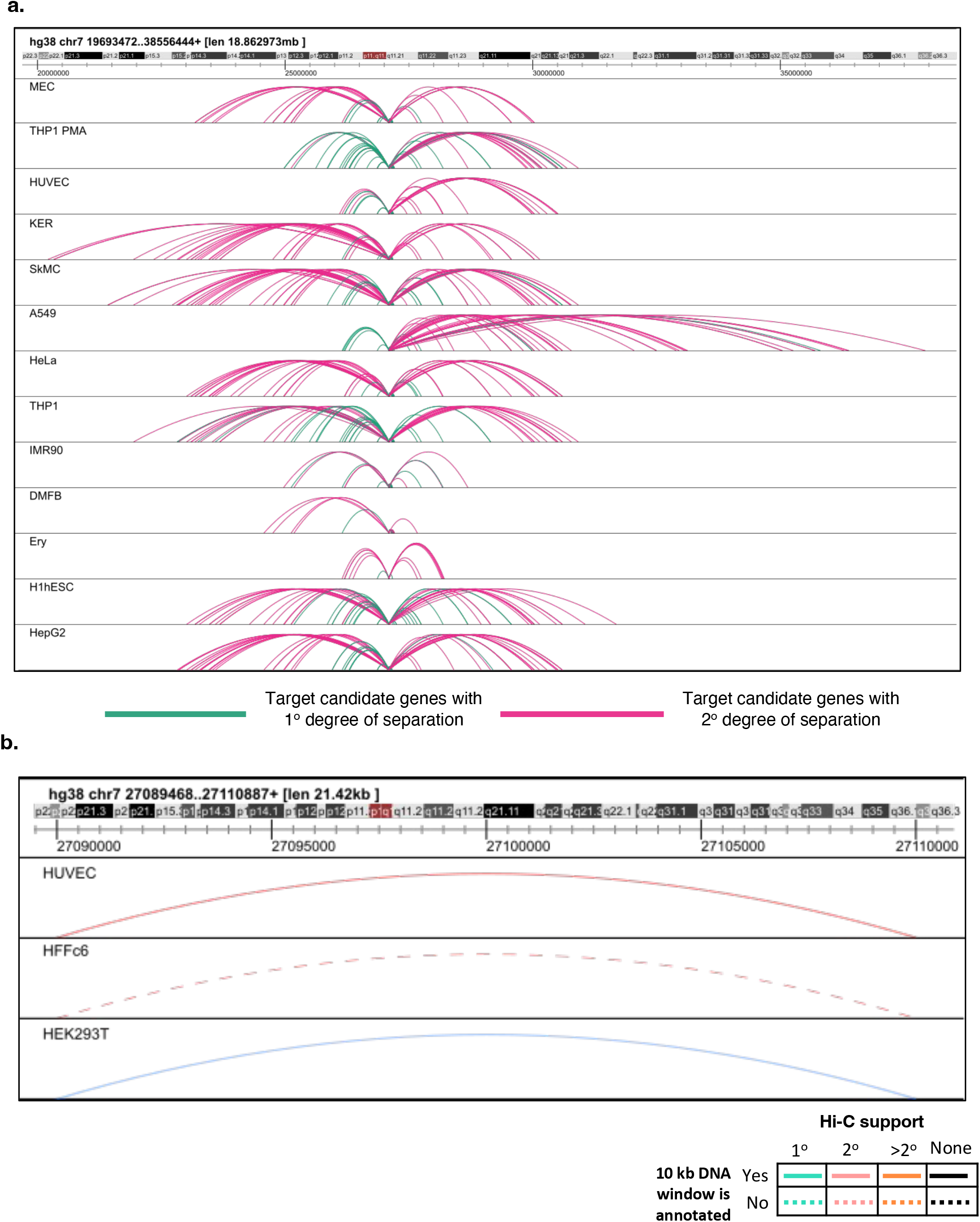
Hi-C and RNA-chromatin interactions for IncRNA. ENSG00000233429. **a.** The top track shows the genomic location of the interaction, followed by tracks showing the Hi-C annotated interactions between lncRNA ENSG00000233429 and its candidate target genes in different cell types. **b.** RNA-chromatin interactions for lncRNA in different cell types. The colors of interactions in cell types HUVEC and HFFc6 tracks show the level of support of RNA-chromatin interactions by Hi-C interactions.

